# Infant Inhibited Temperament in Primates Predicts Adult Behavior, is Heritable, and is Associated with Anxiety-Relevant Genetic Variation

**DOI:** 10.1101/2020.10.30.361477

**Authors:** AS Fox, RA Harris, L Del Rosso, M Raveendran, S Kamboj, EL Kinnally, JP Capitanio, J Rogers

## Abstract

An anxious or inhibited temperament (IT) early in life is a major risk factor for the later development of stress-related psychopathology. Starting in infancy, nonhuman primates, like humans, begin to reveal their temperament when exposed to novel situations. Here, in Study 1 we demonstrate this infant IT predicts adult behavior. Specifically, in over 600 monkeys, we found that individuals scored as inhibited during infancy were more likely to refuse treats offered by potentially-threatening human experimenters as adults. In Study 2, using a sample of over 4000 monkeys from a large multi-generational family pedigree, we demonstrate that infant IT is partially heritable. The data revealed infant IT to reflect a co-inherited substrate that manifests across multiple latent variables. Finally, in Study 3 we performed whole-genome sequencing in 106 monkeys to identify IT-associated single-nucleotide variations (SNVs). Results demonstrated a genome-wide significant SNV near *CTNNA2*, suggesting a molecular target worthy of additional investigation. Moreover, we observed lower p-values in genes implicated in human association studies of neuroticism and depression. Together, these data demonstrate the utility of our model of infant inhibited temperament in the rhesus monkey to facilitate discovery of genes that are relevant to the long-term inherited risk to develop anxiety and depressive disorders.

## INTRODUCTION

An anxious temperament (AT) or inhibited temperament (IT) early in life is a major risk factor for the later development of anxiety and depressive disorders (1–4). Children with an extreme IT early in life are characterized by increased behavioral inhibition and anxiety in novel and potentially threatening contexts. In humans, IT reflects an inhibited nature or disposition that can define how an individual approaches novel situations throughout their life (5–7). Although IT often emerges during the second year of life around the time that a child develops the ability to behaviorally cope with threat (8,9), it can be preceded by increased reactivity in infancy (5). Temperament is partially inherited, and in human samples the heritability estimates range from 20-50% (10–13), (14–16). Because IT is heritable, and a risk-factor for the later development of stress-related psychopathology, IT likely reflects an underlying genetic disposition and/or mediates the relationship between genetic variation and psychopathology.

To understand which molecular components contribute to anxiety and depressive disorders, researchers have begun to perform genome-wide association studies (GWAS) with dispositional anxiety and psychopathology in humans. Large-scale efforts for anxiety disorders (17,18), depressive disorders (19,20), and neuroticism (21) are all ongoing. Polygenic risk-scores and genetic correlation analyses, suggest each of these disorders partially overlap in their genetic substrates(17,21). Although some studies have identified genes with significant single-nucleotide variants (SNVs; e.g. in *CRHR1, ESR1, NTRK2, etc.*)(17,22,23), the specific SNVs influencing psychopathology may not be the same in individuals with different genetic backgrounds and/or who were exposed to different environmental stressors.

Rhesus macaques provide a powerful and well-validated animal model to study the genetic and molecular bases of early-life dispositional anxiety because of their similarity to humans. Next to humans, rhesus macaques are the most widely geographically distributed primates in the world, thereby exhibiting outstanding ecological flexibility and adaptability (24,25). As Old World monkeys, rhesus macaques are phylogenetically close to humans (with a common ancestor ~10-12 million years closer to humans than marmosets and ~50 million years closer than mice). This recent evolutionary divergence between rhesus macaques and humans is reflected in similarities in their genomes, transcript expression, brain circuits, and resulting socio-emotional behavior (26–29). Thus, the rhesus monkey model provides a unique opportunity for translational research that supports and extends the insights gained from human genetic studies.

Capitanio and colleagues have developed and implemented a rhesus monkey model of infant IT, as part of their standardized BioBehavioral Assessment (BBA) that has been applied to over 4000 animals. Infant IT is characterized by below-average expression of latent variables which emerged from behavioral observations spread over two days, termed “Activity” and “Emotionality”, both of which are indicative of behavioral inhibition (30)(see methods for details). Notably, molecular studies of dispositional anxiety in macaques are concordant with human GWAS studies, hinting at evolutionarily conserved mechanisms (e.g. CRH (31,32), and neurotrophic pathways(4,33,34)). Infant IT among macaques provides an excellent model for the study of the human risk for anxiety and depressive disorders (4,35).

The rhesus monkey model of IT provides numerous advantages that facilitate the identification of genes and molecules that contribute to the risk for psychopathology. Rhesus populations have higher levels of genetic variation and lower linkage disequilibrium than do equivalent sample sizes from human populations, likely due to the historic population bottleneck experienced by ancient human populations that gave rise to modern humans (27,36–39). Importantly, this genetic variation is demonstrated in the 853 rhesus macaque whole-genome sequences that Dr. Rogers and colleagues have collected from US primate centers, which has identified >85 million SNVs, including 408,496 missense variants, 9,921 stop codons gained and 7,918 splice acceptor or donor variants (Warren et al. under review). Moreover, the breeding strategy at primate centers results in large multi-generational pedigrees in which each animal has many close and more distant relatives, and this will enrich the population for putatively “rare” variation, thus increasing statistical power in heritability and genetic associations analyses (40).

Here, we demonstrate that infant IT in macaques can be used as an endophenotype to complement human studies of psychopathology, and gain insight into the nature of inherited anxiety. We demonstrate that infant IT is associated with lasting behavioral changes (study 1); show that infant IT is heritable (study 2); and begin to study the genetic variation that underlie this early-life risk-factor (study 3).

## METHOD

### Methods Overview

All studies were performed in rhesus macaques (*Macaca mulatta*) in accordance with the federal guidelines of animal use and care and with the approval of the University of California, Davis Institutional Animal Care and Use Committee. Primary analyses across all studies included animals that underwent BBA testing during infancy (3-4 months of age). Subsets of animals were selected for Food Retrieval Task testing in Study 1 (n=679, 59M/620F), , heritability analyses in Study 2 (n=4433; 2019M/2414F), and whole-genome sequencing in Study 3 (n=106 49M/57F). The only animals that did not undergo BBA-testing were a subset of female animals that underwent test-retest analyses in Study 1 (>2 tests: n=649; >3 tests: n=288; 4 tests: n=88). Additional detailed methods can be found in the *Supplemental Methods* as well as previous publications (30).

### Assessment of Infant IT (Studies 1-3)

The methods for scoring IT in the BBA program have been previously described(30,41). Animals (90-120 days old) are relocated for a 25hr testing period (see supplemental methods). IT is defined based on 4 factors that emerged from factor analyses in a subset of several hundred animals(30). Animals were considered to be “inhibited” if their scores were below the mean for both “Activity” and “Emotionality” across two days of testing, otherwise they were classified as “not inhibited.” The “Activity” factor includes time locomoting; time NOT hanging from the top or side of the cage; rate of environmental exploration; and whether or not the animals ate food*, drank water*, or were seen crouching in the cage* (*=dichotomized due to rarity). The Emotionality factor includes rate of cooing; rate of barking; and whether the animals scratched*, displayed threats*, or lipsmacked* (*=dichotomized due to rarity).

### Food Retrieval Task

The Food Retrieval task was administered at approximately 6:00 am on the day after relocation, prior to morning health and husbandry, by a technician that was blind to infant IT scores. To ensure there was no familiarity with the testing context, the Food Retrieval Task was performed in a different location than BBA testing. The technician stood in front of the cage, then approached the animal and hand presented a food treat for five seconds, taking care to avert her eyes from the monkey (a stopwatch with an audible beep was used for timing). Treat retrieval was recorded. If the animal did not accept the treat, the technician placed the treat on the forage board and stepped back from the cage, averted her eyes, waited five seconds, and again recorded whether or not the treat was retrieved. Three trials were run consecutively for each animal. Because humans in such close proximity can be perceived as threatening, the Food Retrieval task sets up a potential conflict for the animals between a fear of the human versus attraction to a favored treat.

To estimate the relationship between infant IT and refusal to take food in the food retrieval task, we used logistic regressions. To estimate test-retest stability of the dichotomous reach variable, we used chi-squre tests (42,43). All statistical analyses were implemented in Python (version 3.7.3), statsmodels (version 0.10.0; https://www.statsmodels.org/stable/index.html; (44)) was used for regression analyses, and (Pingoin; https://pingouin-stats.org/; (45) was used to perform chi-squared tests.

### Heritability of IT

For this study, we analyzed variation among 4433 infants assessed through the BBA program between May of 2001 and January of 2017. The total sample consisted of 407 inhibited females, 403 inhibited males, 2007 non-inhibited females, and 1616 non-inhibited males. As in our previous work (46–49), all heritability and co-heritability estimates were performed using SOLAR-Eclipse (http://solar-eclipse-genetics.org/). Prior to heritability estimation phenotype variables were normalized using an inverse normal transformation. All heritability analyses controlled for sex. Because all animals were assessed between 3-4 months of age, we did not include Age or Age-squared as covariates in heritability analyses.

### Methods for Whole Genome Sequencing and Mapping

Blood samples were collected from 36 inhibited animals and 70 non-inhibited animals. DNA was extracted and sequenced at the Human Genome Sequencing Center, Baylor College of Medicine using either the Illumina HiSeq 2000 or Illumina HiSeq X Ten system. WGS sequence data for the 106 animals are publicly available through the NCBI SRA (https://www.ncbi.nlm.nih.gov/biosample/?term=Bio+Behavior+Assessment). Paired end reads were aligned to the rhesus Mmul_10 reference using BWA mem with an average mapped sequence depth of 33.66X across the samples. The GATK v. 4.1.2.0 (50) pipeline was used to identify single nucleotide variants (SNVs) and insertions/deletions (indels) smaller than 7bp. Variant Effect Predictor (VEP) (51) was used to annotate variants based on merged Ensembl and RefSeq gene models.

IT-related variants were analyzed using FaST-LMM (52) which implements a linear mixed model that takes potential relatedness into account, controlling for sex. Sequence variants of interest were further examined by lifting the rhesus positions over to the orthologous human position and performing CADD analysis (53).

Permutation tests were used to compare IT-related associations to relevant gene lists extracted from published human genome-wide gene-association studies (GWGAS). First, the minimum p-value for each gene was computed. Then the average minimum p-value for IT-associations for each gene in the target-list was computed and stored. Finally, for each analysis we performed 10,000 permutations with a similarly sized set of randomly selected genes, and determined the average p-value of those gene-sets. The p-value was computed as the proportion of permutations that resulted in a lower p-value than the target gene-set.

## RESULTS

### Study 1: Infant IT predicts later-life behavior

To begin, we determined whether infant IT reflects an animal’s stable disposition throughout their lifespan. We assessed behavioral inhibition during a Food Retrieval Task, when adult animals (n=679) were offered a treat by a human experimenter. We hypothesized that inhibited animals would forgo reward in the presence of this potentially threatening human. Logistic regressions demonstrated animals with high levels of inhibited temperament at 3-4 months of age are less willing to take a treat years later (z=3.248, p=0.001; Fig 1).

**Figure 1.**
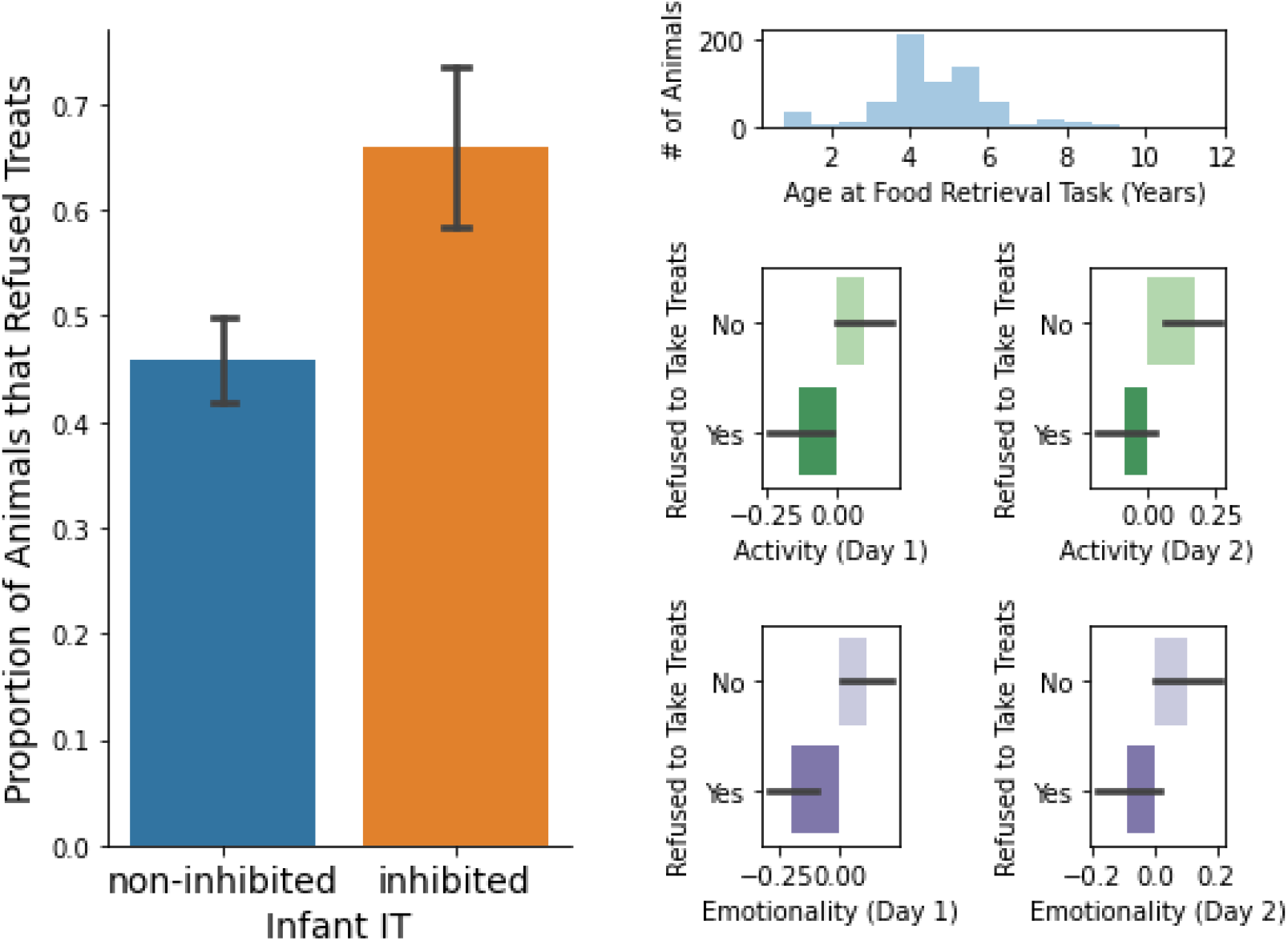
Infant Inhibited Temperament (IT) are more likely to refuse treats from a potentially-threatening human intruder. A) Proportion of animals that refused treats in the Food Retrieval Task in relation to infant IT. B) A histogram of ages for animals when tested in the Treat Refusal Task, on average >4 years after assessment of infant IT. C) The factors that contribute to IT, i.e. Day 1 and 2 Activity and Emotionality, are each related to the propensity to refuse treats.

IT is a composite measure of 2 latent factors, across 2 days, i.e. Day 1 and Day 2 Activity & Emotionality. To be sure each factor of IT was contributing to the underlying temperamental variable, we assessed each variable as a predictor of treat refusal. Analyses revealed each measure, Activity and Emotionality, on each day to be predictive of treat refusal (all p’s < 0.014). Supplementary analyses revealed no significant effects of sex (p=.209); significant effects of IT on treat refusal in both males (t=2.710, p=0.007) and females (t=3.193, p=0.001); an effect of age on treat refusal, such that older animals were less likely to refuse a treat (z=-5.419, p<.001); and that IT remained significantly associated with treat refusal while controlling for age (z=3.785, p<.001) (see *Supplemental Results* for additional descriptions). Interestingly, although animals have multiple opportunities to retrieve treats, the Food Retrieval Task typically results in an all-or-nothing result, with only ~17% (49/292) of animals who did not retrieve a treat on the first trial going on to retrieve any treat. Unsurprisingly, we obtained similar results when examining first-treat refusal, for IT (z=3.248 p=0.001), age (z=-6.064, p<.001), and IT controlling for age (2.935, p=0.003). Together these data suggest that IT as assessed in this protocol is a trait-like measure, which is susceptible to change with experience, but does remain detectably consistent within an animal across contexts as they mature.

We next evaluated treat-refusal as a stable assessment. In a separate group of 649 female animals, we performed the food retrieval task multiple times, longitudinally. Animals received 2 (N=649) to 4 (N=88) assessments, on average 1.8 years apart. Chi-squared tests revealed treat-inhibition to be stable between each assessment (p’s<.001; Fig 2), such that 69% of animals responded consistently during the 4th assessment on average 5.24 years later (range=1.9-9.2 years). Again, treat refusal tended to be all-or-nothing, with 82% of animals that refused the first treat refusing all treats (chi^2^=430.68, p<.000001), and first-treat retrieval showing similar stability (p’s<.001).

**Figure 2.**
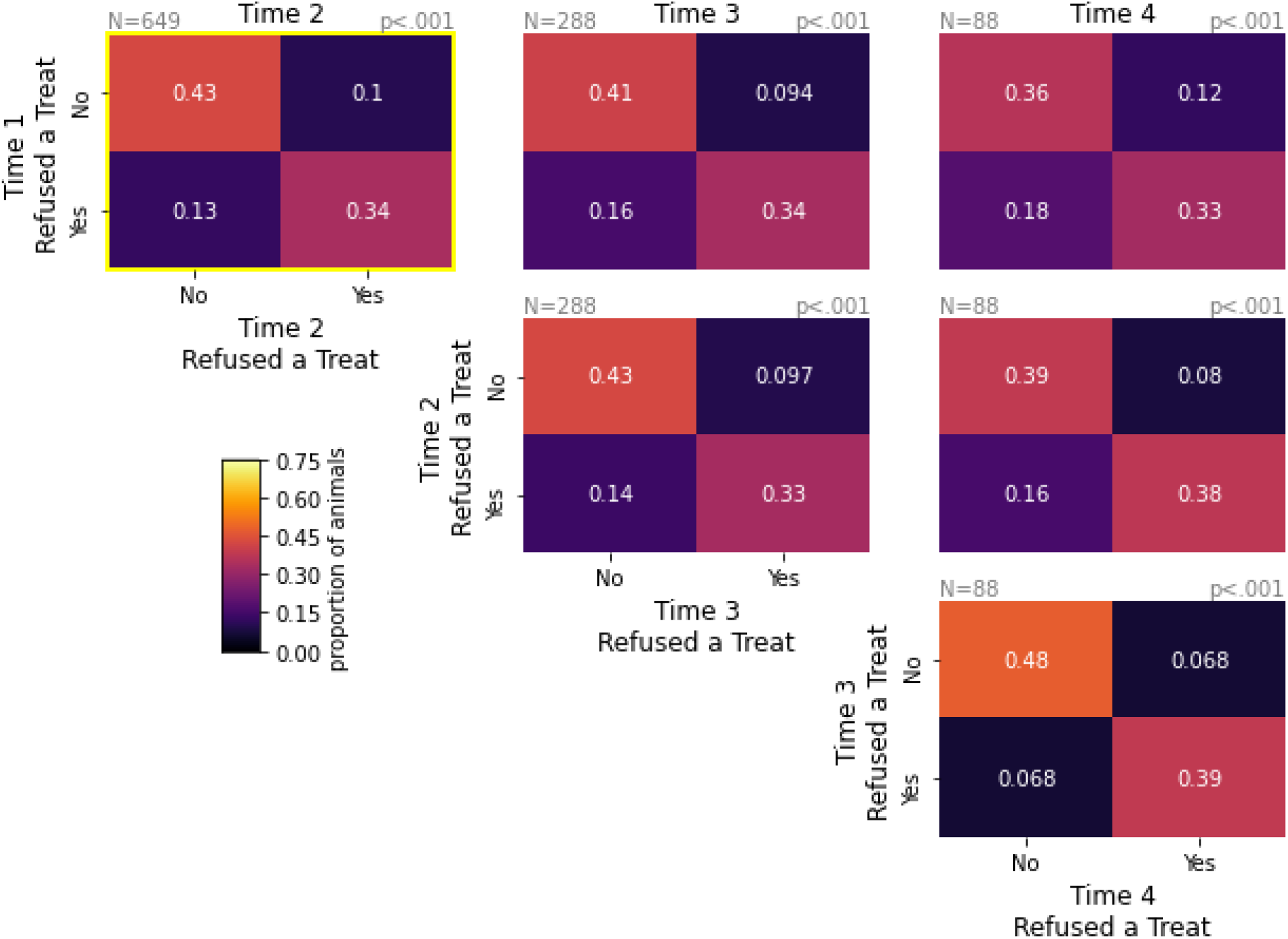
Treat refusal in the Food Retrieval Task is stable over multiple assessments. Contingency tables with the proportion of animals that refused a treat across assessments. The N and p-value for the chi-squared test are above each contingency table in grey.

Together, these data demonstrate infant IT reflects life-long behavioral inhibition, that reflects varied behaviors across novel contexts.

### Study 2: Infant IT is heritable

We next examined the heritability of IT, and its contributing factors, across 4433 animals that were part of a large multi-generational pedigree. Results showed that IT is significantly heritable with h^2^=0.19 (std_h2_=0.027; p=3.7e-27; Fig 3a). Results also found IT’s latent factors, i.e. Day 1 and Day 2 Activity and Emotionality, to be significantly heritable (p’s < 1.0e-16) with h^2^ values ranging from 0.17-0.30. To test the extent to which the latent factors contributing to IT resulted from overlapping genetic variation, we performed genetic correlation analyses. Genetic correlation analyses test the extent to which covariation in phenotypes are associated with covariation in estimated genetic variation, as estimated by relatedness. Results demonstrated all latent factors that comprise IT were significantly genetically correlated with each other (rho-g estimates from 0.45-0.89; p’s< 0.0001) and with IT (rho-g estimates 0.65-0.90, p’s<1×10^−9^; Fig 3b). These data suggest a partially shared genetic substrate that contributes to these latent factors.

**Figure 3.**
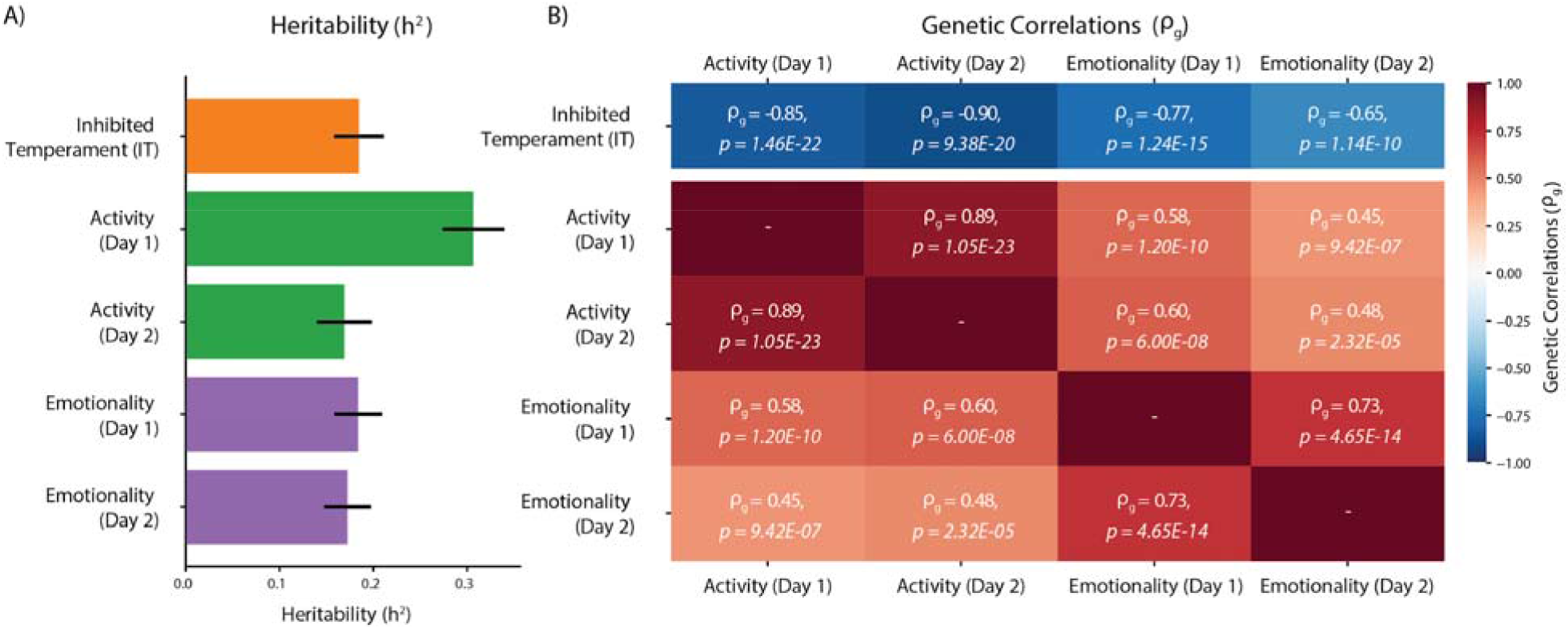
Inhibited Temperament (IT) and its latent factors are heritable, and genetically correlated. Inhibited Temperament (IT) is heritable, as are the contributing latent factors Activity and Emotionality across both days of testing (A). In addition, IT shows a significant negative genetic correlation with each of latent factors, which in turn are positively correlated with each other.

**Figure 4.**
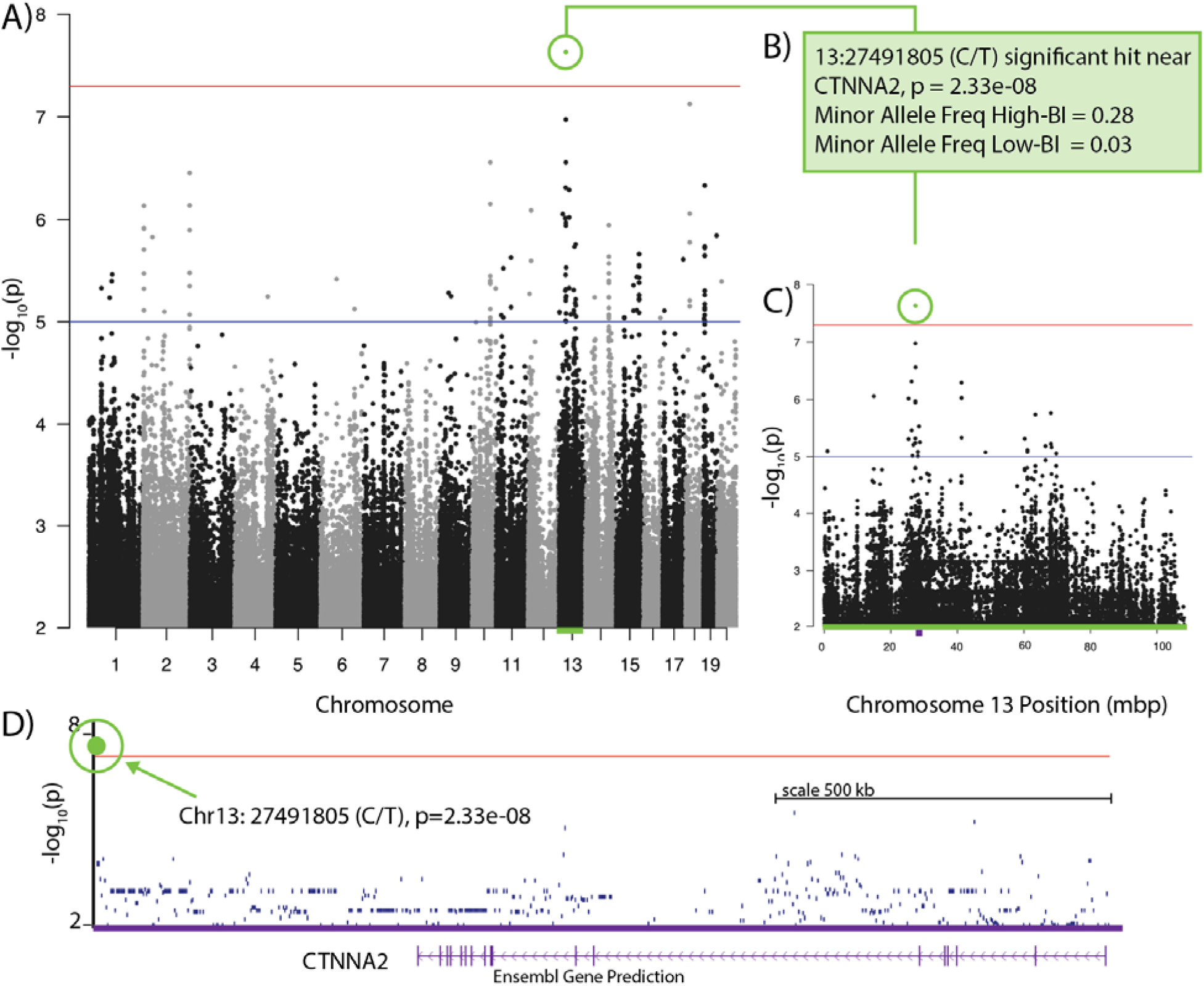
A SNV on Chromosome 13 near *CTNNA2* is associated with IT. A) Genome-wide Manhattan plot of BioBehavioral Assessment (BBA) associations with SNVs. The x axis is the chromosomes and y-axis is the −log_10_ of the FaST-LMM p-value. The genome-wide significance threshold p < 5×10^−8^ is represented by the red line. The significant SNV is highlighted in green. B) A description of the significant SNV. C) Chromosome 13 Manhattan plot of BioBehavioral Assessment (BBA) associations with SNVs. The x axis is the chromosomes and y-axis is the −log_10_ of the FaST-LMM p-value. The genome-wide significance threshold p < 5×10^−8^ is represented by the red line. The significant SNV is highlighted in green. D) UCSC Genome Browser view of SNVs with FaST-LMM −log_10_ p-values in the vicinity of *CTNNA2*. The genome-wide significance threshold p < 5×10^−^8 is represented by the red line.

Together, the data from Study 2 demonstrate infant IT is significantly heritable, making it an ideal starting point to identify genetic variation. Here, we focus genetic analyses on IT which is likely to identify genetic variation that contribute to multiple latent variables, and are less likely to be specifically related to an individual feature of inhibited temperament (e.g. propensity toward locomotion).

### Study 3: GWAS of IT

In this study, we surveyed BBA subjects for DNA sequence polymorphisms that might reasonably be hypothesized to influence IT, and determined whether the results were consistent with relevant human GWAS studies. Across the 106 individuals, we identified 53,030,128 SNVs and 6,435,882 indels using whole genome sequencing, and used FaST-LMM to identify IT-related SNVs, taking relatedness into account. We identified a single SNV (13:27491805:C:T) that exceeded a significance threshold p < 5×10^−8^ which is a standard threshold for GWAS genome wide significance (Table S1, Fig. 5). No indels met this threshold. Two additional nearby SNVs had the third and fourth lowest p-values in the dataset, but failed to reach the genome wide significance threshold (Table S1, Fig. 5B), including 13:27444729:G:C, 47,076bp upstream and 13:27493293:T:A, which was 1477bp downstream of the top SNV. All of these SNVs are intergenic, but 13:27491805:C:T is 473,868 bp downstream of the 3’ end of Catenin Alpha-2 (*CTNNA2*). However, although follow-up analyses revealed 4 indels and 43 SNVs in *CTNNA2*, we were unable to identify variation that was both related to IT (p<0.05) and predicted to be functional (i.e. CADD>10, see supplementary results).

**Figure 5.**
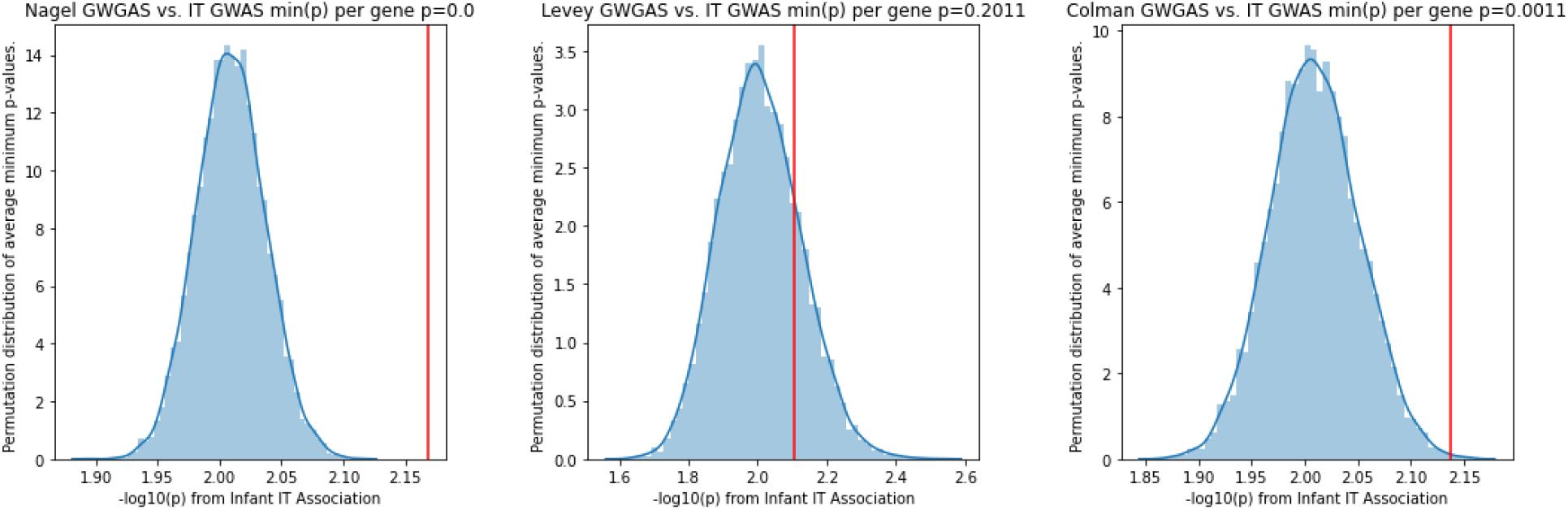
Permutation analyses of the minimum p-value for each gene from human GWGAS studies. Permutation tests revealed the average minimum p-value from genes found in human GWGAS studies of Neuroticism (p<0.001) and Major Depressive Disorder (p=0.011) was significantly lower than the average minimum p-value in equally sized random subsets of genes. This same analysis did not reach significance for a GWGAS of Anxiety Disorders (p=0.201).

In addition to our genome-wide significant hit in *CTNNA2*, we relaxed the formal genome-wide significance threshold and explored variation in other genes that were marginally significantly (p<.01) associated with IT (Table S4), but do not reach genome-wide significance. Interestingly, our results revealed uncorrected associations with genetic variation in genes we’ve previously implicated in inhibition, including an 3’ UTR variant in *NTRK3* (p=0.005) and an intronic variant in *PRKCD* (p=.007) (33,34,54). Such associations must be considered less meaningful but can provide indications of genetic effects that deserve further study.

Using permutation analyses, we tested for IT-related enrichment in genes relevant to stress-related psychopathology to determine if further genetic studies of monkey IT are likely to identify evolutionarily-conserved genes and pathways that are relevant to human psychopathology. Because species differences prevent base-pair by base-pair comparisons, we examined gene-level enrichment based on genome-wide gene-association studies (GWGAS) from studies of human Neuroticism (547 genes; Negal)(21), Anxiety Disorders (31 genes; Levey)(17), and Depressive Disorders (251 genes; (23). We examined the minimum IT-related p-value for target genes deemed significant in published GWGAS studies. Permutation analyses found the average minimum p-value to be significantly lower in target gene lists as compared to randomly selected genes, for neuroticism (Negal: p<.0001) and depressive disorders (Colman: p=.0018), but not Anxiety Disorders (Levey: p=.21; though this list only contained 31 genes) (Fig 6). Interestingly, genes that were significant in the human GWGASs and the current analyses (p<.005, uncorrected) include *CTNNA2*, *PRKCD*, *NTRK3* and *ESR1*, highlighting the potential for identifying evolutionarily conserved mechanisms. These data highlight the promise of our nonhuman primate approach, and suggest that additional data will identify additional IT-related genetic signals.

## DISCUSSION

An early-life inhibited temperament represents one of the strongest known risk-factors for the later development of psychopathology. Here, we demonstrate infant IT in rhesus macaques to be a heritable endophenotype that reflects stable context-independent behavior, and demonstrates promise for identifying molecules that contribute to stress-related psychopathology in humans. The data suggest infant IT reflects underlying biological processes that manifest across different behaviors in different contexts. These data mirror the work done in humans, in which children classified as behaviorally inhibited tend to remain inhibited as they mature, and are at a substantial risk to develop stress-related psychopathology (1–3,55–59).

The extremely large number of phenotyped animals (N>4000) that are part of a large multi-generational pedigree, provides compelling evidence for heritability, and a unique opportunity for further study of the specific SNVs that contribute to IT. Though generally consistent with our previous behavioral genetics studies of adolescent/adult rhesus macaques (35,37,47,60–63), previous heritability estimates tended to be slightly higher (e.g. ~25%-40% heritable: (46–49,60). The analyses we describe here include over four-thousand animals and found infant IT to be ~19% heritable (see Fig 3). It is possible that these heritability estimates will increase as animals mature, as has been observed in twin studies of human anxiety(64,65). Like with human anxiety and depressive disorders, infant IT was associated with the family structure, such that inhibited animals are more likely to be related to inhibited animals.

Our preliminary genome-wide association analyses revealed a promising hit near the *CTNNA2* gene in chromosome 13, which encodes Catenin alpha-2. This is of particular interest because *CTNNA2* has been previously implicated in human association studies, including a GWGAS of anxiety disorders, and encodes a neuron-specific catenin that is important for cell-to-cell adhesion and synaptic plasticity, is expressed throughout cortical and subcortical structures, and is implicated in numerous studies of IT-related phenotypes (17,66–73). The association analysis presented here does not constitute evidence for a definitive association between *CTNNA2* and infant IT in the rhesus monkey. That said, together with findings in humans, these data contribute to the rationale for further study of *CTNNA2* in anxiety-related behavior in animal models, and highlight a potential molecular mechanism that may drive stable anxiety across the lifespan. More generally, our results are consistent with many molecules contributing to IT, reflecting disruption across many of the brain regions and cell-types that are thought to play a role in anxiety and threat-processing (See supplemental discussion) (31,33,34,47,48,54).

Moreover, these data suggest that further study of infant ITs genetics may identify molecules that are relevant to human psychopathology that can be targeted for mechanistic study. In this sample of 106 rhesus macaques, we identified IT-associations that were enriched for genes identified in GWGAS studies of ~449k subjects phenotype for neuroticism and ~624k subjects phenotyped for major depressive disorder in humans (including ~185k patients). We did not find significant enrichment for the genes implicated in a GWGAS of ~200k subjects phenotyped for anxiety disorders. This is likely because this study only identified 31 genes and will likely increase as the number of anxiety disorder genes goes up. Interestingly, although not significantly enriched for average p-value, the *CTNNA2* gene was identified in the million veterans analysis of anxiety, suggesting another relevant gene, like CRHR1(31), is shared between human anxiety disorders and rhesus monkey infant-IT. Together, these data highlight the utility of using our rhesus monkey model of infant IT to identify molecules that may be relevant to human psychopathology, and suggest there is a shared molecular substrate that is conserved across primates.

### Conclusion

Elucidating the genes and genetic mechanisms that predispose individuals to stress-related psychopathology can inform our understanding of psychopathology, and guide the development of biologically-informed treatments. Understanding of disease mechanisms and developing optimal treatments requires human epidemiological insights, as well as well-validated animal models. With the power afforded by a pedigree-based whole-genome sequencing approach in a genetically variable species, nonhuman primate studies may be able to identify genes and specific genetic variants that would otherwise take hundreds of thousand of subjects using a traditional GWAS approach (27,39,40,74). Here, we help provide support for a translational model that can support and extend the insights gained from human genetic studies to identify the mechanisms that underlie stress-related psychopathology.

## Acknowledgements

We would like to thank the staff at the California National Primate Research Center, Harsha Doddapaneni, Donna Muzny and the HGSC data production team, Dr. William Mason for helping to develop the Food Retrieval Task, as well as H. Sompura. This work was supported by NIH grants OD010962 to JPC, R01MH121735 to ASF, OD011173 to JR, and the California National Primate Research Center (P51OD011107).

## Author Contributions

**Fox AS:** Conceptualization, Formal Analysis, Methodology, Software, Visualization, Writing – original draft (lead); **Harris RA:** Data curation, Formal Analysis, Software, Visualization, Writing – review & editing; **L Del Rosso:** Conceptualization (Food Retrieval), Investigation (BBA and Food Retrieval) (lead), Project Administration (BBA program) (lead), Supervision (BBA) (lead), Writing – review and editing; **M Raveendran:** Data curation, Formal analysis, Project administration; **Kamboj S:** Writing – original draft; **EL Kinnally:** Resources (provided DNA), Writing-review and editing **Capitanio JP:** Conceptualization (BBA program), Data curation (BBA), Methodology (BBA), Resources (BBA data), Writing – review and editing; **Rogers J:** Conceptualization, Supervision, Writing – review and editing.

## Supplemental Materials

### Supplemental Methods

#### Subjects: Overview

All subjects were rhesus macaques (*Macaca mulatta*). Primary analyses across all studies included animals that underwent BBA testing during infancy (3-4 months of age). Subsets of animals were selected for Food Retrieval Task testing in Study 1 (n=679; 59M/620F), heritability analyses in Study 2 (n=4433; 2019M/2414F), and whole-genome sequencing in Study 3 (n=106; 49M/57F). The only animals that did not undergo BBA-testing were a subset of female animals that underwent test-retest analyses (n=649F) in Study 1. Details on each group can be found below. All studies were performed in accordance with the federal guidelines of animal use and care and with the approval of the University of California, Davis Institutional Animal Care and Use Committee.

#### Assessment of Infant IT (Studies 1-3)

Across all studies, inhibited temperament status in infant rhesus macaques was identified using the California National Primate Research Center’s (CNPRC) BioBehavioral Assessment (BBA) program. This program, which has been described in detail elsewhere (30,35) comprises a series of standardized behavioral and physiological assessments, conducted over a 25-hr long period when infant macaques ranged from 90-120 days of age. Assessed animals were from all four rearing environments at CNPRC: field cages, corn cribs, indoor mother reared colony, or nursery reared colony. Field cages enclosures (approximately half an acre each) house approximately 50–200 animals each. Corn cribs are roughly 400 square foot outdoor enclosures that house approximately 15–30 animals each. Field cage and corn crib reared animals were raised in their respective outdoor enclosure with their biological or foster mother. Indoor mother reared and nursery reared animals were raised indoors with no social group. Indoor mother reared animals were housed in a cage with their biological or foster mother, and at most one additional adult female and infant macaque pair. Nursery reared animals were individually housed until 3 weeks of age, at which time they were given visual access to an infant of the same age, and eventually paired with this peer at 5 weeks.

Subjects are separated from their mothers, arrive in the testing area at 0900, and are housed in standard laboratory cages (0.58 × 0.66 × 0.81 m, Lab Products, Maywood, NJ). Animals are tested in cohorts of 5-8 animals. Beginning at 0915h on Day 1, a trained observer records the behavior of each animal for five minutes using a predetermined random order and a standard ethogram for this species (see Golub et al., 2009). This assessment is repeated at 0700h the next day (Day 2). At 1000h on Day 2, infants are returned to their mothers, who had been housed in a separate area of the facility, and allowed to nurse for an hour before all animals are returned to their home cages.

As described by Golub et al. (2009) (30), behavioral data from these assessments were subjected to an exploratory factor analysis using a sample of several hundred animals. Once a factor structure was identified, a confirmatory factor analysis, involving several hundred different animals was conducted to confirm the structure. These data revealed a two factor structure that was replicable: Activity (proportion of time spent locomoting; proportion of time spent not hanging from the side of the cage, rate of environmental exploration, and dichotomous codes for whether the animal ate food, drank water, or crouched) and Emotionality (rate of cooing, rate of barking, and dichotomous codes for whether the animal scratched, displayed threats, or lipsmacked) (definitions of all behaviors are found in Golub et al. (2009) (30). For each year, four scales were created, Activity and Emotionality for Day 1, and separately for Day 2, and values for each were z-scored with mean of 0 and standard deviation of 1. Animals were considered “inhibited” if their scores were below the mean for all four scales, and were otherwise classified as “not inhibited.”

##### Study 1

###### Subjects: IT and Food Retrieval task (Study 1a)

The subjects were 679 (n=59 male) primarily female macaques that were relocated from an outdoor enclosure (either field cages or corn cribs) for various reasons (project harvest, shipping, cage relocation etc). No animals were paired. Animals were relocated by early afternoon, and the Food Retrieval test was administered at approximately 0600h the next day, prior to morning health and husbandry. This approach minimizes any effects of separation and relocation on the animals, by leveraging separations that are a part of normal animal and veterinary care.

###### Subjects: Test-Retest of Food Retrieval task (Study 1b)

Test-retest was done on a separate group of adult female animals, many of whom did not undergo infant BBA testing. Subjects were 649 female subjects that underwent multiple food retrieval tests (at least 2 tests: n=649; at least 3 tests: n=288; 4 tests: n=88). The method for repeated assessment was identical to the initial testing.

###### Food Retrieval Task

The Food Retrieval task was administered at approximately 6:00 am on the day following the animal’s relocation, and prior to morning health and husbandry, by a technician that was blind to infant IT scores. In all cases, the Food Retrieval Task was performed in a different location than BBA testing, to ensure there was no familiarity with the testing context. The technician stood in front of the animal’s cage, then approached the animal and hand presented a food treat (Condition 1) for a period of five seconds, taking care to avert her eyes from the monkey (a stopwatch with an audible beep was used for timing). If the treat was taken during the five seconds, this was recorded. Any behavioral responses directed toward the technician were also recorded (data not shown). If the animal did not accept the treat, the technician took a step back, recorded the animal’s position in the cage, stepped forward, placed the treat on the forage board and stepped back from the cage (Condition 2), averted her eyes, waited five seconds, and again recorded whether or not the treat was retrieved as well as any behavioral response directed toward the technician. Three such trials were run consecutively for each animal. Because humans in such close proximity can be perceived as threatening, the Food Retrieval task sets up a potential conflict for the animals between a fear of the human versus attraction to a favored treat.

###### Statistical Analyses: Food Retrieval Task

To estimate the relationship between infant AT and refusal to take food in the food retrieval task, we used logistic regressions. To estimate test-retest stability of the dichotomous reach variable, we used chi-squre tests(42,43). All statistical analyses were implemented in Python (version 3.7.3), statsmodels (version 0.10.0; https://www.statsmodels.org/stable/index.html;(44)) was used for regression analyses, and (Pingoin; https://pingouin-stats.org/; (45)) was used to perform chi-squared tests.

##### Study 2

###### Subjects: Heritability of IT

For this study, we analyzed variation among 4433 infants (3-4 months of age) that were assessed for infant-IT as a part of the BBA program between May of 2001 and January of 2017. Parentage was assessed through breeding, blood typing, DNA. Since 1999 parentage was established with microsatellite markers(75,76); before that, a combination of approaches were taken, including blood typing and breeding records. The total sample consisted of 407 inhibited females, 403 inhibited males, 2007 non-inhibited females, and 1616 non-inhibited males.

###### Statistical Analyses: Heritability Estimation

All heritability and bivariate heritability analyses were performed using SOLAR-Eclipse (http://solar-eclipse-genetics.org/). In brief, heritability is estimated using maximum likelihood, as the proportion of total of variance in an N-by-N pair-wise phenotype-covariance matrix based on the relatedness between animals (77,78). Prior to heritability estimation phenotype variables were normalized using an inverse normal transformation. All heritability analyses controlled for sex. Because all animals were assessed between 3-4 months of age, we did not include Age or Age-squared as covariates in heritability analyses.

Genetic correlations (◻_) were estimated using bivariate heritability analyses in SOLAR-Eclipse(47,79,80). Bivariate heritability analyses are performed using methods similar to the heritability analyses detailed above, with a covariance matrix that represents both traits and their interaction. Significance was determined by comparing the full model to a second model in which the ◻_◻_ parameter was fixed to be 0.

##### Study 3

###### Subjects: Whole-Genome-Sequencing

A total of 106 animals assessed for infant IT, 36 inhibited animals and 70 non-inhibited animals, were selected for whole-genome sequencing (n=49 male, n=57 female). Animal selection was based on the following criteria: field cage born and reared; reared by biological mother (ie, foster-reared animals excluded); minimally related (kinship coefficient < 0.03125); DNA available in our bank (minimal DNA for animals born in 2005-2008, so those animals were excluded).

###### Methods for genome sequencing and mapping

Rhesus macaque DNA samples (n=106) provided by Dr. John Capitanio and Dr. Erin Kinnally were sequenced at the Human Genome Sequencing Center, Baylor College of Medicine using either the Illumina HiSeq 2000 or Illumina HiSeq X Ten system. WGS sequence data for the 106 animals are publicly available through the NCBI SRA (https://www.ncbi.nlm.nih.gov/biosample/?term=Bio+Behavior+Assessment). The paired end reads were aligned to the rhesus Mmul_10 reference genome assembly using BWA mem with an average mapped sequence depth of 33.66X across the samples. The GATK v. 4.1.2.0 (50) pipeline was used to identify single nucleotide variants (SNVs) and insertions/deletions (indels) smaller than 7bp. Variant Effect Predictor (VEP) (51) was used to annotate variants based on merged Ensembl and RefSeq gene models.

###### Statistical Analyses: Genome-Wide Associations

Variants were analyzed for association using FaST-LMM (52) which implements a linear mixed model that takes into account potential relatedness among samples. FaST-LMM was run using the BioBehavioral Assessment (BBA) of inhibited (n=36) or not inhibited (n=70) as the phenotype and sex as a covariate.

Sequence variants of interest were further examined by lifting the rhesus positions over to the orthologous human position and performing CADD (53) analysis which predicts the functional impact of variants. CADD integrates annotations from multiple genomic resources and calculates a score that predicts the functional impact of a variant. A CADD score of 10 indicates that the variant is predicted to be among the 10% most functional variants across the entire genome. A CADD score of 20 means that the variant falls in the top 1% most functional.

###### Statistical Analyses: Permutation Analyses to Compare with Published Genome-wide gene-association studies (GWGAS)

We compared our results to 3 genome-wide gene-association studies (GWGAS) on human Neuroticism (547 genes; (21), Anxiety Disorders (31 genes; (17), and Depressive Disorders (251 genes; (23)). A list of relevant genes was extracted from each published GWGAS study. To perform the permutation analysis, we first computed the minimum p-value for each gene. We computed the average minimum p-value in our IT-association, in the genes reported in each list. Then, for each analysis we performed 10,000 permutations with a similarly sized set of random genes, and determined the average p-value of those gene-sets. The p-value was computed as the proportion of permutations that resulted in a lower p-value than the target gene-set.

### Supplemental Results

#### Exploratory analyses for IT and behavioral inhibition in the Food Retrieval Task

Exploratory logistic regressions were performed to examine potentially confounding variables of sex or age during Food Retrieval Task. There were no significant effects of sex (p=.209). Additionally, IT was significantly associated with treat refusal in both males (t=2.710, p=0.007) and females (t=3.193, p=0.001), separately. There was substantial variation in the age at which animals were exposed to the Food Retrieval Test (Fig 1). Logistic regressions found that age was significantly correlated with treat refusal, such that older animals were less likely to refuse a treat (z=-5.419, p<.001). Importantly, IT remained significantly associated with treat refusal when entered simultaneously with age in a logistic regression (z=3.785, p<.001).

Interestingly, although animals have multiple opportunities to retrieve treats, the Food Retrieval Task typically results in an all-or-nothing, result, with only ~17% (49/292) of animals who did not retrieve a treat on the first trial going on to retrieve any treat. Unsurprisingly, our main results were maintained when restricting Food Retrieval Task data to refusal of the first treat, for IT (z=3.248 p=0.001), age (z=-6.064, p<.001), and IT controlling for age (2.935, p=0.003). Together these data suggest that IT as assessed in this protocol is a trait-like measure, which is susceptible to change with experience, but does remain detectably consistent within an animal across contexts as they grow up.

#### Genome-wide significant hit and CTNNA2

We searched for predicted functional variants by reciprocally lifting over each SNV to the orthologous human position and annotating each SNV with CADD information including the calculated CADD PHRED Score (Table S2). The genome-wide significant hit, 13:27491805:C:T is in an open chromatin region based on Ensembl annotations. This could have regulatory implications, but the low CADD score suggests it may not. The nearby SNV 13:27444729:G:C is annotated as intergenic, but it does have a higher CADD score (CADD=4.436). The human nucleotide orthologous to 13:27493293:T:A could not be identified with existing information and tools.

Because the variant exhibiting genome-wide significant association with IT was near *CTNNA2*, we further examined SNVs and indels in that gene without imposing a FaST-LMM p-value threshold. There were 4 indels in the UTRs of *CTNNA2*, but the most significant FaST-LMM p-value was 0.14. There were 43 SNVs in *CTNNA2* including 28 UTR variants, 14 synonymous variants and a single missense variant. The 13:28743267:C:T missense variant (Table S3) has a high CADD score of 28.5 suggesting it may have functional significance, but the p-value (p=0.48) suggests it is unlikely to have an association with IT, and the allele frequency difference between inhibited and non-inhibited animals is small. Thus, the genome-wide significant result near *CTNNA2* seems to affect IT via an unknown mechanism.

### Supplementary Discussion

Our preliminary genome-wide association analyses revealed a promising hit near the *CTNNA2* gene in chromosome 13. *CTNNA2* encodes catenin alpha-2 (also called cadherin-associated protein, alpha 2). Catenin alpha-2 is a neuron-specific catenin that is important for cell-to-cell adhesion and synaptic plasticity, and is expressed throughout cortical and subcortical structures (brain-map.org). Using a pedigree-based analysis in humans, researchers showed that biallelic loss of *CTNNA2* results in severe deficits in neuronal migration accompanied by intellectual impairment and autism-like features (73). The only study to our knowledge to examine the mechanistic role of *CTNNA2* in fear/anxiety-related behavior found that *CTNNA2* knockout mice have a deficiency in fear-potentiated startle (66). Despite the limited understanding surrounding the function of *CTNNA2* in anxiety, numerous discovery-based analyses point to its relevance for psychopathology. Most excitingly Levey et al., (2019)(17) found SNPs in human CTNNA2 to be associated with anxiety disorders in their GWGAS (1/31 genes tested above). Consistent with a shared genetic substrate between anxiety, depression, and addiction, GWAS studies in humans have also implicated variation in the *CTNNA2* gene in impulsivity (67), excitement seeking (68), depression with comorbid substance abuse (69), substance dependence (70), anorexia(71), and chronic pain (72). As noted in the main manuscript, the association analysis presented here does not constitute evidence for a definitive association between *CTNNA2* and infant IT in the rhesus monkey. That said, together with findings in humans, these data contribute to the rationale for further study of *CTNNA2* in anxiety-related behavior in animal models, and highlight a potential molecular mechanism that may drive stable anxiety across the lifespan.

In addition to our finding near the *CTNNA2* gene, the data point to the potential involvement of other molecular systems in infant IT. Studies in humans and nonhuman primates have identified an evolutionarily conserved distributed neural circuit that underlies anxiety and anxiety-like behavior. Importantly, this circuit includes cortical regions (e.g. orbital proviso cortex and insular regions [OPro/AI)], extended amygdala regions (e.g. central nucleus of the amygdala [Ce], and bed nucleus of the stria terminalis [BST)], as well as regions of the brainstem (e.g. periaqueductal gray [PAG)]. In nonhuman primates, we have demonstrated that metabolism and connectivity across these circuits is heritable, providing a neural substrate that can mediate the effects of genes on anxiety(47,48). We predict that the heritability and genetic effects are mediated by alterations within these circuits. For example, though it did not survive multiple comparison correction, we identified IT-related variation in *ESR1* and *NTRK2*, which have been implicated in human GWGAS studies, as well as *NTRK3* and *PRKCD*, which have been mechanistically implicated in anxiety-like behavior in animal models. These findings should be interpreted cautiously, as they did not reach formal levels of significance, but suggest a possible convergence with human genetics studies and rhesus RNA-sequencing studies that should be explored with additional statistical power.

Consistent with the results presented here, mechanistic studies in nonhuman primates and rodents have implicated specific molecules and cell-types within these regions that contribute to these distributed alterations in anxiety-related brain function. For example, we have demonstrated that expression of *NTRK3* is associated with anxiety-related responding (33,34). Moreover, experimental manipulation of NTRK3-signalling in the dorsal amygdala region is sufficient to increase anxiety-related behavior in adolescent rhesus macaques(34). The data presented here, hint at additional support for the role of *NTRK3* in the risk to develop anxiety and depressive disorders by implicating a 3’ variant as potentially contributing to infant IT. Along similar lines, rodent studies have identified cells expressing PKC∂ in the lateral Ce (CeL) that, when optogenetically stimulated, result in decreased freezing to a cue (81). Moreover, these neurons seem to be required for fear learning in the BLA(82), and required for the effects of benzodiazepines(83). These data are supported by RNA-sequencing studies in the nonhuman primate where Kovner and colleagues identified expression of the gene that encodes PKC∂, *PRKCD*, to be associated with behavioral inhibition in response to potential threat (54). Interestingly, Kovner et al., demonstrated Ce PKC∂-expressing cells project to BST, suggesting a potential mechanism for the heritability of Ce-BST functional connectivity(49,54). That said, again, these findings should be interpreted cautiously, and considered as modest evidence worthy of further investigation, rather than proof of these specific SNVs or genes as relevant to IT.

There are likely thousands of genes(84) and near infinite SNVs that contribute to anxiety and depressive disorders. Identifying any single gene that is causally related to a phenotype implicitly implicates related molecules in the disorder. For example, when an individual non-synonymous SNV implicates a particular G-protein coupled receptor, this suggests the molecules that bind that receptor, regulate that receptor, or mediate its intracellular effects also may play a role in that disorder. For example, identifying BI^+^-related polymorphism in the intracellular loop of the *CRHR1* gene(31) implicates CRH ligand (32), and intracellular signaling kinases that interact with CRHR1 in the expression of anxiety. The cascade of implicated molecules does not stop there. Genes involved in causing CRH to be released and other receptors on CRH-expressing cells can also be considered likely to play a role in the risk to develop stress-related psychopathology. Although each of these implicitly related molecules is unlikely to be the cause of psychopathology in any given patient, that does not mean that they cannot be useful for treatment targets.

## Supplementary Tables

**Table S1.** SNVs of interest based on FaST-LMM p-value including the allele nucleotides and alternate allele frequency by BBA status.

| SNV ID | Ref Allele | Alt Allele | FaST-LMM p-value | Inhibited Alt AF | Not inhibited Alt AF |
| --- | --- | --- | --- | --- | --- |
| 13:27491805:C:T | C | T | 2.33e-08 | 0.2778 | 0.02899 |
| 13:27444729:G:C | G | C | 1.06e-07 | 0.4028 | 0.08696 |
| 13:27493293:T:A | T | A | 2.76e-07 | 0.2571 | 0.02899 |

**Table S2.** SNVs lifted over to the human genome and CADD results

| SNV ID | Human (hg38) | Ensembl Consequence | CADD Phred Score |
| --- | --- | --- | --- |
| 13:27491805:C:T | 2:81106490 | Regulatory – Open Chromatin | 0.344 |
| 13:27444729:G:C | 2:81148697 | Intergenic | 4.436 |

**Table S3.** *CTNNA2* functional SNVs

| SNV ID | Consequence | Ref Allele | Alt Allele | CADD Phred Score | FaST-LMM p-value | Inhibited Alt AF | Not inhibited Alt AF |
| --- | --- | --- | --- | --- | --- | --- | --- |
| 13:28743267:C:T | missense | C | T | 28.5 | 0.48 | 0 | 0.0072 |

## Notes

### Competing Interest Statement

The authors have declared no competing interest.

